# OpusTaxa: A Unified Workflow for Taxonomic Profiling, Assembly, and Functional Analysis of Shotgun Metagenomes

**DOI:** 10.64898/2026.04.15.718825

**Authors:** Yen-Kai Chen, Clarice M. Harker, Cong M. Pham, Luke Grundy, Hannah R. Wardill, Michael J. Roach, Feargal J. Ryan

## Abstract

Shotgun metagenomics has become a cornerstone of microbiome research, yet the complexity of existing workflows remains a major barrier for life scientists without dedicated bioinformatics support. Manual database setup, detailed sample sheet preparation, and management of software dependencies can make routine analysis difficult and time-consuming. Cross-study comparisons are further hampered by inconsistent processing pipelines, database versions, and profiling strategies, limiting reproducibility and the potential for large-scale meta-analyses. We present OpusTaxa, an open-source Snakemake workflow that provides end-to-end processing of short paired-end shotgun metagenomic data with minimal configuration. Users provide either FASTQ files or Sequence Read Archive accessions; OpusTaxa automatically downloads required databases, performs quality control, removes host reads, and executes taxonomic profiling, metagenome assembly, and functional analysis. All analysis modules can be independently toggled, and per-sample outputs are automatically merged into harmonised, cross-sample tables ready for downstream exploration. Across two public datasets, we demonstrate how OpusTaxa can be used to compare consistency across complementary taxonomic profilers and to estimate microbial load in addition to standard metagenomic workflows.

**Availability:** OpusTaxa is freely available at https://github.com/yenkaiC/OpusTaxa. Documentation, test data, and example configurations are included in the repository.

## Introduction

The global microbiome community has generated hundreds of thousands of metagenomes in recent decades. Yet differences in processing pipelines, database versions, and profiling strategies make cross-study comparisons difficult and often irreproducible. As bioinformatic methods and reference databases evolve rapidly, even datasets from the past few years can yield new insights when re-analysed with updated tools. Similarly, as analysis pipelines differ in reference databases, underlying algorithms, and reporting conventions, results can be method-dependent [1], complicating comparisons across studies. At the same time, running multiple complementary tools on the same dataset can provide an internal sanity check, helping to distinguish stable biological signals from tool-specific artefacts. However, re-analysing past data with up-to-date databases and software versions is currently a time-consuming and complex process. Workflow management systems such as Snakemake [2] and Nextflow [3] allow users to simplify this process. While there are existing workflows for metagenome analyses (**Table 1**), they focus primarily on one of taxonomic profiling [4] or metagenome assembly [5-7], and often require users to manually install tools and download external databases [8, 9]. Additionally, few tools automate integration with data repositories to support reanalysis of public data [5].

**Table 1:**
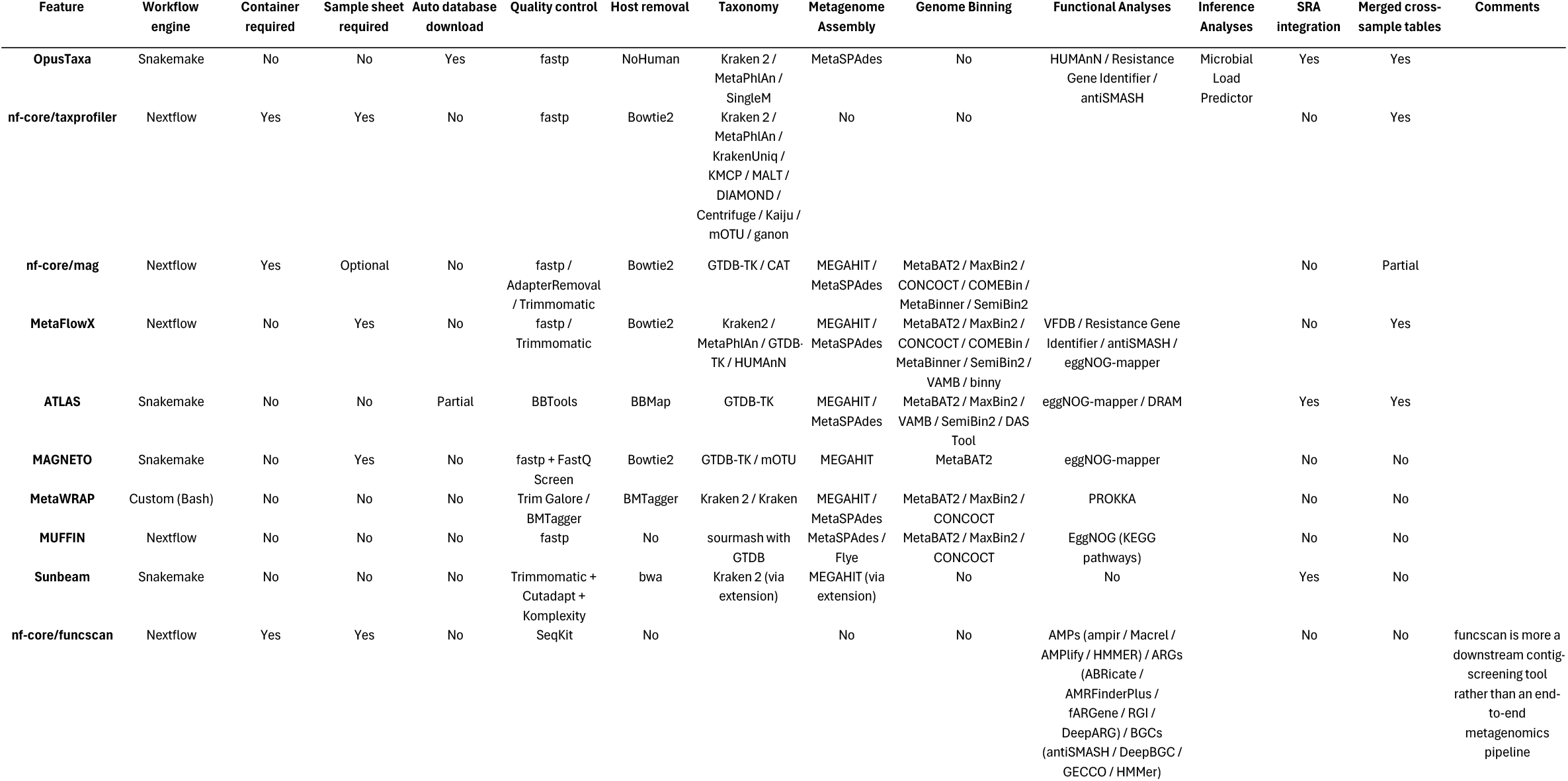
Feature comparison across publicly available workflows for shotgun metagenomic data analysis. Workflows were evaluated for workflow engine, container and sample sheet requirements, automated database provisioning, quality control and host removal strategies, taxonomic profiling tools, metagenome assembly, genome binning, functional and inference analyses, SRA integration, and generation of merged cross-sample output tables.

To fill this gap, we developed OpusTaxa, an open-source Snakemake workflow designed to provide end-to-end metagenomic processing for life scientists with minimal bioinformatics expertise.

### Design philosophy and architecture

OpusTaxa is designed to be a comprehensive and easy-to-use metagenomic profiling workflow manager for short-read paired-end shotgun metagenomic sequencing data, such as that generated by Illumina and MGI platforms (**Figure 1**). It is built on Snakemake [2] and offers both conda and containerisation for reproducible dependency management. The workflow is modular: every analysis tool can be toggled on or off via command (e.g. metaphlan=true, kraken2=false), allowing users to customise their analysis without modifying pipeline code.

**Figure 1:**
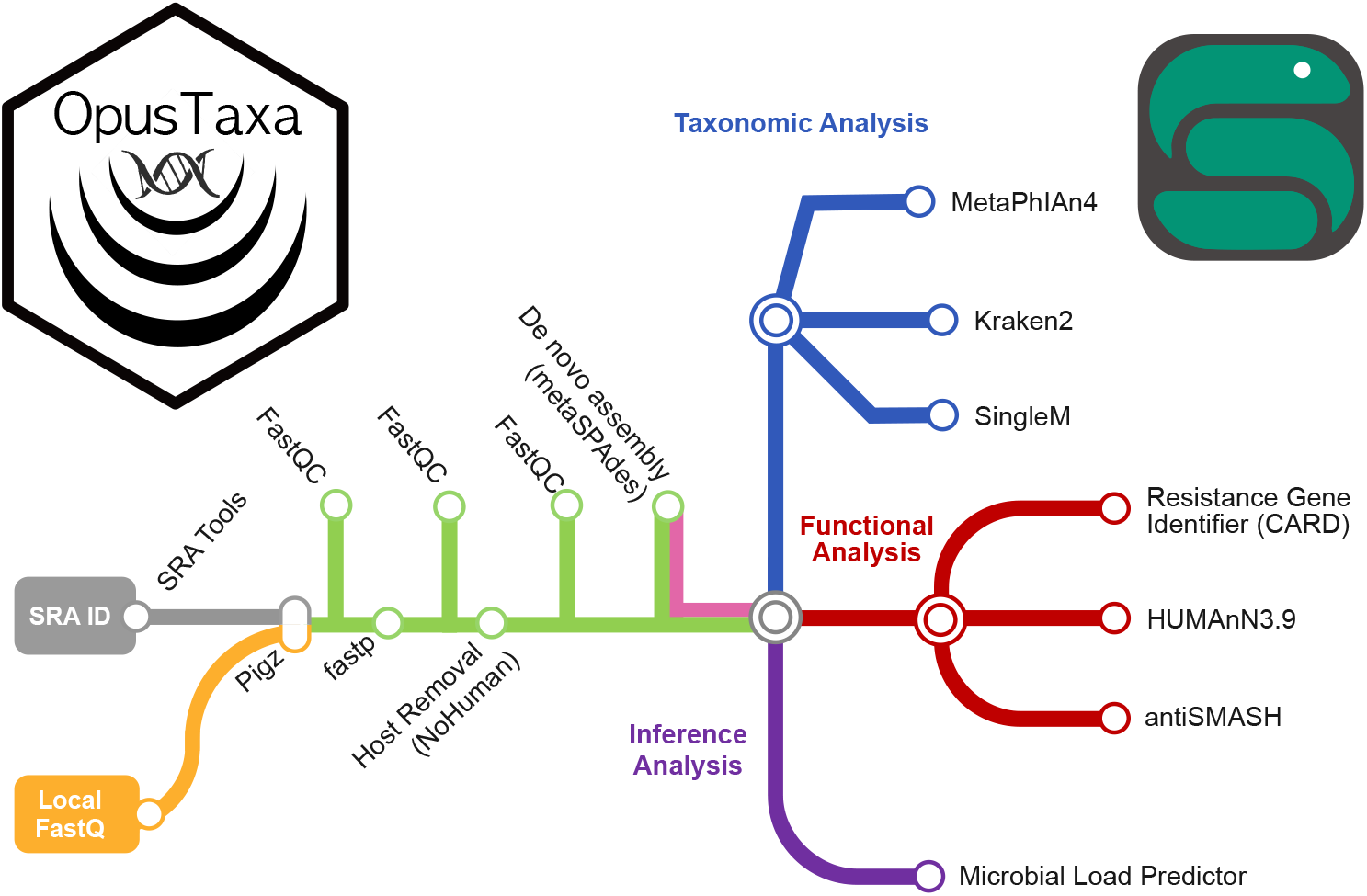
Pipeline overview of OpusTaxa. Modules are colour-coded by analysis category: data acquisition, quality control, host removal, taxonomic profiling, metagenome assembly, and functional analysis.

A key design goal was to minimise the configuration burden. OpusTaxa automatically detects paired-end FASTQ files in the input directory, supporting common naming conventions (Illumina, SRA, dot-separated). All required databases for selected tools including those for NoHuman [10], MetaPhlAn [11], SingleM [12], Kraken2 [13], HUMAnN [14], antiSMASH [15], and CARD [16] are provisioned automatically on first run via checkpoints. Snakemake’s built-in job tracking ensures that previously completed samples are not reprocessed when the pipeline is re-run with additional data. For high-performance computing environments, OpusTaxa ships with the option of using a Snakemake profile for SLURM job submission, with configurable resource allocation per rule (CPU, memory, runtime). Default resource settings are tuned for typical metagenomic datasets but can be overridden in the configuration.

### Input format and Sequence Read Archive integration

OpusTaxa can either process local FASTQ files or download data directly from the Sequence Read Archive (SRA) [17] via fasterq-dump. Downloaded reads are compressed with pigz [18] and automatically standardised from SRA to Illumina-style filenames. Local and SRA-derived samples can be mixed in a single run, making it straightforward to reanalyse published cohorts alongside new data.

### Quality Control and host-read removal

Raw paired-end reads are quality-trimmed with fastp [19], which removes adapters, low-quality bases, and short fragments. Quality reports are generated at three stages, raw reads, post-trimming, and post-host-removal, using FastQC [20] and aggregated with MultiQC [21] to provide a clear view of data quality at each step. This three-stage reporting gives users a clear view of data quality at each processing step, enabling rapid identification of problematic samples. Human reads are removed using NoHuman [10], which employs a Kraken2-based classification against the Human Pangenome Reference Consortium (HPRC) r2 database. This approach offers an optimal host-read removal while maintaining computational efficiency, as pangenome databases better capture human genomic diversity than a single linear reference genome.

### Taxonomic profiling tools

OpusTaxa currently integrates three complementary taxonomic profilers that differ in methodology and output. MetaPhlAn 4 [11] identifies species using clade-specific marker genes, producing relative abundance profiles. SingleM [12] takes a distinct approach, using single-copy marker genes built from the Genome Taxonomy Database to estimate both taxonomic composition and the prokaryotic fraction of the metagenome [22]. Kraken2 [13], paired with Bracken [23] for Bayesian abundance re-estimation, offers k-mer-based classification of each individual read, which includes archaea, bacteria, viruses, fungi, protozoa, and the human genome. These tools were selected to provide complementary taxonomic profiles based on distinct analytical strategies and reference databases, enabling users to compare results across fundamentally different approaches. Each profiler can be enabled independently, and OpusTaxa automatically merges per-sample profiles for each tool into cross-sample abundance tables for easy downstream analysis and comparison between samples.

### De-novo assembly and Functional Profiling

When enabled, OpusTaxa performs per-sample metagenome assembly with MetaSPAdes [24], retaining contigs and scaffolds and removing intermediate files to conserve storage. OpusTaxa supports both read-based and contig-based functional characterisation and automatically merges results across samples. For reads, HUMAnN 3.9 [14] quantifies gene families and metabolic pathways using UniRef90 and outputs are normalised to copies-per-million. As HUMAnN does not natively accept paired-end input, it is run on forward reads only. For contigs, Resistance Gene Identifier (CARD) [16] annotates antimicrobial resistance genes, while antiSMASH [15] detects biosynthetic gene clusters to infer secondary-metabolite potential. To improve runtime and the quality of the output, contigs under 1000 base pairs are not included in functional analyses. OpusTaxa also estimates absolute microbial load from MetaPhlAn output profiles via Microbial Load Predictor [25], facilitating quantitative microbiome profiling. All per-sample outputs are consolidated into harmonised, cross-sample tables optimised for comparative analyses.

OpusTaxa is developed in accordance with the FAIR Principles for Research Software (FAIR4RS) [26]. The source code is openly available on GitHub under MIT licence, with documentation, test data, and example configurations included in the repository. A versioned, citable release is archived on Zenodo (DOI: https://doi.org/10.5281/zenodo.19491844), providing a persistent identifier for reproducibility. Dependencies are fully specified via conda environment files, and all outputs conform to standard bioinformatics file formats to facilitate interoperability with downstream tools.

## Results

To demonstrate OpusTaxa’s utility, we first re-processed four publicly available human gut metagenome samples from the SRA. The five most abundant species identified in each sample by each of the three taxonomic profilers (Bracken, MetaPhlAn 4, and SingleM) were compared across samples (**Figure 2**). Across all four samples, the three profilers recovered broadly concordant dominant community members, SingleM reported alternative scientific names in a small number of instances (e.g. *Agathobacter faecis* in place of *Roseburia faecis*), reflecting differences in taxonomy frameworks rather than major disagreement in biological interpretation. This concordance is notable because the three profilers operate with different databases and algorithms. Bracken, derived from Kraken2, uses k-mer-based classification of individual reads against a broad reference database; MetaPhlAn 4 infers species abundance from clade-specific marker genes; and SingleM uses universal single-copy marker genes derived from the Genome Taxonomy Database. Agreement across these complementary approaches suggests that the dominant signals in these samples are robust to method choice

**Figure 2:**
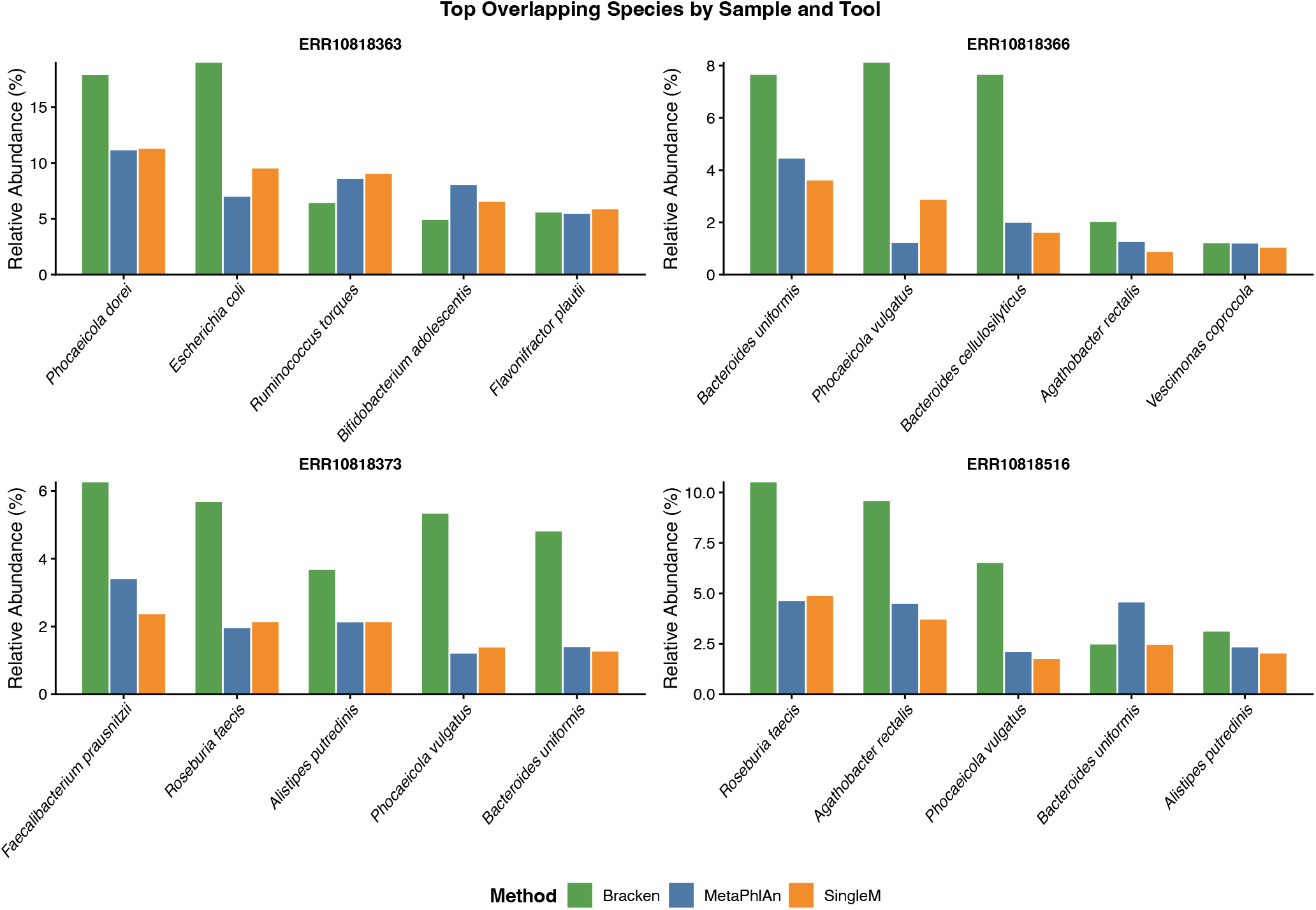
Cross-tool taxonomic concordance across 4 control samples of human gut metagenome reanalysed from Duru et al., 2018 [28]. The five most abundant species identified per sample by Bracken, MetaPhlAn 4, and SingleM are shown. All samples were processed using OpusTaxa v0.3.

We next applied OpusTaxa to a longitudinal dataset examining gut microbiome recovery following treatment with three broad-spectrum antibiotics (gentamicin, meropenem, and vancomycin) [27]. OpusTaxa downloaded the raw reads directly from the SRA, generated species-level abundance profiles with MetaPhlAn 4, and supplied these profiles to the Microbial Load Predictor (MLP). Shannon diversity calculated from MetaPhlAn 4 abundance profiles (**Figure 3a**) revealed a diverse and evenly distributed community at baseline that collapsed sharply following antibiotic administration, with diversity at its lowest on day 4. Partial recovery was observed by day 8, with near recovery at day 42, and complete recovery by day 180, reproducing reported patterns of post-antibiotic microbiome reconstitution [27]. Predicted microbial load (**Figure 3b**) mirrored this trajectory, declining following antibiotic exposure and recovering by day 42, showing similar results to a prior study [25]. Together, these analyses show that OpusTaxa can recover biologically plausible and literature-consistent patterns from public metagenomic datasets while also extending interpretation through integrated quantitative modules such as MLP.

**Figure 3:**
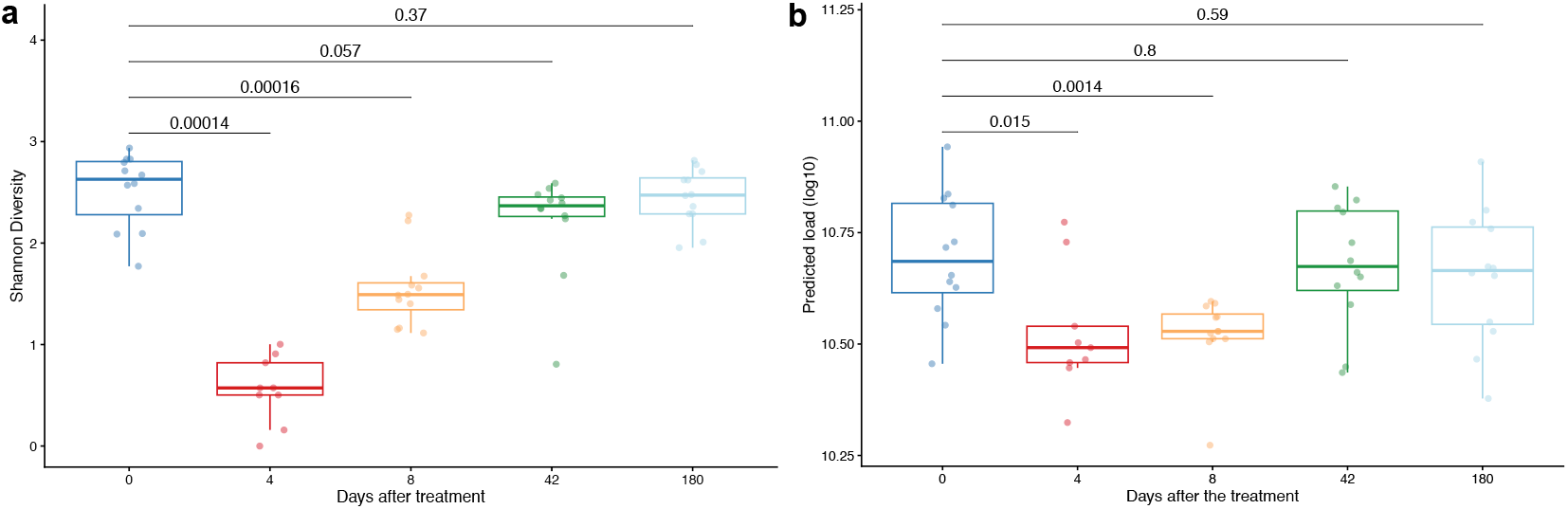
Recovery of gut microbiota following broad-spectrum antibiotic exposure, reanalysed from Palleja et al., 2018). Subjects were treated with gentamicin, meropenem, and vancomycin. (a) Shannon diversity index calculated from MetaPhlAn 4 species-level abundance profiles across sampling timepoints. (b) Predicted microbial load (cells per gram) estimated by Microbial Load Predictor across the same timepoints. All samples were processed using OpusTaxa v0.3. Statistical comparisons between day 0 and each subsequent timepoint (days 4, 8, 42, and 180) were performed using the Wilcoxon rank-sum test.

## Conclusion

OpusTaxa provides an end-to-end Snakemake workflow for shotgun metagenomic analysis that combines taxonomic profiling, metagenome assembly, and functional analysis in a single, minimally configured framework. Its design priorities: automatic database provisioning, no sample sheet requirement, conda-only dependency management, and built-in SRA integration; distinguish it from existing workflows and lower the barrier for life scientists who lack dedicated bioinformatics support. OpusTaxa is in active development with further taxonomic profilers planned (for example mOTU [29] and others in consideration), as well as virome profiling and additional functional profiling tools. Currently, OpusTaxa deliberately omits genome binning, and does not perform taxonomic profiling of assembled contigs, though OpusTaxa’s MetaSPAdes outputs can be used as input for these tools if desired To our knowledge, OpusTaxa is the first published workflow to integrate SingleM and the Microbial Load Predictor directly within an end-to-end shotgun metagenomic processing framework. In summary, OpusTaxa enables rapid, scalable analysis of either public or local metagenomic data with up-to-date analysis tools and databases in as few as five short commands from installation to results.

## Acknowledgements

We thank colleagues for helpful discussions that informed development of OpusTaxa and users for testing. Large language models were utilized to assist with grammar and spelling corrections, as well as for code documentation.

## CRediT authorship contribution statement

**Yen-Kai Chen**: Conceptualization, Software, Formal analysis, Validation, Visualization, Writing original draft. **Clarice M. Harker**: Validation, Writing review & editing. **Cong M. Pham**: Validation, Writing review & editing. **Luke Grundy**: Supervision, Writing review & editing. **Hannah R. Wardill**: Supervision, Writing review & editing. **Michael J. Roach**: Conceptualization, Software, Methodology, Supervision, Writing review & editing. **Feargal J. Ryan**: Conceptualization, Methodology, Supervision, Funding acquisition, Writing review & editing.

